# Effective *in-vitro* inactivation of SARS-CoV-2 by commercially available mouthwashes

**DOI:** 10.1101/2020.12.02.408047

**Authors:** Katherine Davies, Hubert Buczkowski, Stephen R Welch, Nicole Green, Damian Mawer, Neil Woodford, Allen DG Roberts, Peter J Nixon, David W Seymour, Marian J Killip

## Abstract

Infectious SARS-CoV-2 can be recovered from the oral cavities and saliva of COVID-19 patients with potential implications for disease transmission. Reducing viral load in patient saliva using antiviral mouthwashes may therefore have a role as a control measure in limiting virus spread, particularly in dental settings. Here, the efficacy of SARS-CoV-2 inactivation by seven commercially available mouthwashes with a range of active ingredients were evaluated *in vitro*. We demonstrate ≥4.1 to ≥5.5 log_10_ reduction in SARS-CoV-2 titre following a one minute treatment with commercially available mouthwashes containing 0.01-0.02% stabilised hypochlorous acid or 0.58% povidone iodine, and non-specialist mouthwashes with both alcohol-based and alcohol-free formulations designed for home use. In contrast, products containing 1.5% hydrogen peroxide or 0.2% chlorhexidine gluconate were ineffective against SARS-CoV-2 in these tests. This study contributes to the growing body of evidence surrounding virucidal efficacy of mouthwashes/oral rinses against SARS-CoV-2, and has important applications in reducing risk associated with aerosol generating procedures in dentistry and potentially for infection control more widely.

## MAIN TEXT

SARS-CoV-2 is the virus responsible for causing COVID-19 (1), and infectious SARS-CoV-2 is detectable in the oral cavities and the saliva of COVID-19 patients (2, 3) with potential implications for disease transmission. Aerosol-generating procedures, particularly in the dental setting, therefore pose a potential infectious risk to health care teams working in close proximity to patients while these procedures are being carried out (4). The World Health Organization recommends the use of pre-procedural mouth rinses for the reduction of SARS-CoV-2 viral load in patient saliva as a control measure for reduction of this infectious risk (5). Here, we have assessed seven different commercially available mouthwashes with a range of active ingredients for the efficacy against SARS-CoV-2 *in vitro*.

The commercial mouthwashes tested in this study are listed in Table 1. All products were stored in their original packaging according to manufacturer’s instructions and were unopened prior to testing. *In vitro* SARS-CoV-2 inactivation assessments were performed in a containment level 3 facility, and all virus manipulations were performed within a class III microbiological safety cabinet. Briefly, one volume of virus preparation (SARS-CoV-2 England 2 strain, in tissue culture fluid [TCF] comprising Minimum Essential Media [MEM] and 5% foetal calf serum) was mixed with ten volumes of product and mixed well by inversion. Products were incubated at 20°C (+/− 2°C) for one minute, then immediately titrated in phosphate-buffered saline (PBS) to generate a ten-fold dilution series. Dilution series were directly applied to 96-well plates of Vero E6 cells to determine the 50% tissue culture infectious dose (TCID50) as previously described (6). All products were tested in triplicate, and a triplicate set of samples treated with an equivalent volume of PBS was included in each experiment as a control for virus recovery. Mean titre reductions were calculated by subtracting the mean log_10_ titre of treated samples from the mean log_10_ titre of PBS-treated samples. The cytotoxicity of treated samples varied between products, and a cytotoxic control sample comprising one volume of PBS to ten volumes of product was evaluated in parallel and used to calculate the limit of detection for each product.

**Table 1:** SARS-CoV-2 inactivation by commercial mouthwashes.

| Product | Manufacturer | Active ingredient/s<sup>‡</sup> | Mean titre reduction;<br>log<sub>10</sub> TCID<sub>50</sub>/ml (95% CI) |  |
| --- | --- | --- | --- | --- |
|  |  |  | TCF<br>unconcentrated | TCF<br>concentrated |
| Chlorhexidine Gluconate Antiseptic Mouthwash (Peppermint Flavour) | Ecolabs | 0.2% chlorhexidine gluconate (formulation contains ethanol) | 0.5 (0.1-0.9) | Not tested |
| Corsodyl (Alcohol Free Mint Flavour) | GlaxoSmithKline | 0.2% chlorhexidine gluconate (alcohol-free formulation) | 0.4 (-0.2-0.7) | Not tested |
| Listerine Advanced Defence Sensitive | Johnson & Johnson | 1.4% dipotassium oxalate (alcohol-free formulation) | ≥ 3.5** (3.2-3.8) | ≥ 4.2** (3.9-4.4) |
| Listerine Total Care | Johnson & Johnson | Eucalyptol, thymol, menthol, sodium fluoride, zinc fluoride | ≥ 4.1* (3.8-4.4) | ≥ 5.2* (4.9-5.4) |
| OraWize+ | Aqualution Systems | 0.01-0.02% stabilised hypochlorous acid | ≥ 5.5<sup>†</sup> (5.2-5.8) | 0.4 (0.0-0.8) |
| Peroxyl | Colgate | 1.5% hydrogen peroxide | 0.2 (-0.1-0.5) | Not tested |
| Povident | Huddersfield Pharmacy Specials | 0.58% povidone iodine | ≥ 4.1* (3.8-4.4) | ≥ 5.2* (4.9-5.4) |

<sup>‡</sup>Principal active ingredient/s listed by the manufacturer only are given; refer to manufacturer documents for full ingredients <sup>†</sup>Limit of detection was 0.7 log<sub>10</sub> TCID<sub>50</sub>/ml <sup>\*</sup>Limit of detection was 1.7 log<sub>10</sub> TCID<sub>50</sub>/ml due to product cytotoxicity <sup>\*\*</sup>Limit of detection was 2.7 log<sub>10</sub> TCID<sub>50</sub>/ml due to product cytotoxicity

Two Listerine compositions were evaluated in this study: Listerine Advanced Defence Sensitive and alcohol-free Listerine Total Care. Both formulations reduced SARS-CoV-2 titre to below the limit of detection for the tests after a one minute treatment: ≥3.5 log_10_ reduction for Listerine Advanced Defence Sensitive and ≥4.1 log_10_ reduction for Listerine Total Care, respectively (Table 1). The high level of cytotoxicity associated with Listerine Advanced Defence Sensitive meant that the reduction we could demonstrate for this product in this test was below the >4 log_10_ reduction given in the standard for virucidal quantitative suspension tests, BS EN 14476 (7). Previously, we have conducted a wide range of chemical inactivation testing to inform risk assessments around sample processing for the COVID-19 response (6, 8); we have used purification methods extensively for these assessments to remove components that are cytotoxic in cell culture and would otherwise increase the limit of detection for treated samples. However, we have found these methods unsuitable for evaluation of short (e.g. two minutes or less) treatment times due to the additional time required for sample processing. To see if we could increase the detectable titre reduction without performing a post-treatment purification step, we tested these products using a concentrated virus preparation, generated by concentrating TCF containing virus through 100-kDa-cutoff Amicon Ultra-15 centrifugal filters. When tested against this concentrated virus, we could demonstrate ≥4.2 log_10_ titre reduction for Listerine Advanced Defence Sensitive and ≥5.2 log_10_ for Listerine Total Care. Both of these products were therefore clearly effective at inactivating SARS-CoV-2 in a TCF matrix, despite both products differing in their active ingredients. The manufacturer lists 1.4% dipotassium oxalate as the active ingredient in Listerine Advanced Defence Sensitive, while eucalyptol, thymol, menthol, sodium fluoride and zinc fluoride are given as active ingredients for Listerine Total Care, although the contribution of these particular ingredients to the antiviral activity of these mouthwashes is unclear. Alternative Listerine compositions have been evaluated for SARS-CoV-2 antiviral activity by others, including Listerine Cool Mint (9, 10), Listerine Antiseptic (11) and Listerine Advanced Gum Treatment (10). This study provides evidence that Listerine Advanced Defence Sensitive and Total Care formulations are similarly effective against SARS-CoV-2.

Povident contains 0.58% povidone iodine, and reduced SARS-CoV-2 titre by ≥4.1 log_10_ in our tests using unconcentrated TCF and ≥5.2 log_10_ using concentrated TCF (Table 1). This is consistent with previous studies of povidone iodine-based products, where efficacy in vitro against coronaviruses has been demonstrated, including against SARS-CoV-1 and Middle East respiratory syndrome-associated coronavirus MERS-CoV (12, 13). More recently, oral rinse products containing between 0.5% and 1.0% povidone iodine have been demonstrated to be effective against SARS-CoV-2 in vitro (9, 10, 14, 15) and in reducing viral load in the saliva of human COVID-19 patients (16).

OraWize+, a product containing 0.01-0.02% hypochlorous acid (HOCl) as its active ingredient, reduced virus titre in unconcentrated TCF by ≥5.5 log_10_ TCID50/ml, to below the limit of detection for the assay (Table 1). A potential role for hypochlorous acid-based products as oral rinses to combat SARS-CoV-2 has been proposed (17, 18), but to our knowledge this is the first *in vitro* evidence for efficacy of a hypochlorous acid-based mouthwash against SARS-CoV-2. It is important to note however that OraWize+ was not effective when tested against concentrated TCF (Table 1), potentially due to high levels of protein in this sample matrix, suggesting that the chemistry of this product may be affected by complex samples types. This is an observation we have also made for other hypochlorous acid-based inactivants (unpublished data) and further testing is required to determine the significance of this observation for product use.

Two chlorhexidine gluconate-based products were evaluated in this study: Corsodyl (alcohol-free) and Ecolabs Chlorhexidine Gluconate Antiseptic Wash (containing ethanol). Neither were effective at inactivating SARS-CoV-2 (Table 1), consistent with previous studies demonstrating only a very small effect on SARS-CoV-2 (9, 10). Peroxyl (containing 1.5% hydrogen peroxide) was similarly ineffective. This last observation was initially surprising considering that one minute treatment with 0.5% hydrogen peroxide has been reported to be effective against human coronavirus 229E in virus suspension tests (19) and that 1% hydrogen peroxide pre-procedural mouth rinse is recommended by the World Health Organisation (WHO) and others for reduction of infectious risks in the context of COVID-19 (4, 5). However, ours is not the only study to demonstrate minimal in vitro effectiveness of hydrogen peroxide-based mouth rinses against SARS-CoV-2 and the superior effectiveness of other types of oral rinses (9, 15).

The availability and stability of these products vary, and these factors may impact their utility in different settings. OraWize+ has a much shorter shelf life than other products tested (one month after opening) and must be protected from light; we have found that it can lose effectiveness when stored incorrectly (unpublished data). Povident has a relatively short shelf life, and is not widely available in the UK (indeed, currently there is no widely commercially available povidone iodine mouthwash in the UK). In contrast, the Listerine formulations tested have a considerably longer shelf life, are far more widely available and are designed for use by the general public.

In conclusion, we have demonstrated effective inactivation of SARS-CoV-2 by by Listerine Advanced Defence Sensitive and Total Care formulations, and by commercial mouthwashes containing 0.01-0.02% hypochlorous acid or 0.58% povidone iodine in *in vitro* tests using TCF. Our data support the use of these products, but not the use of hydrogen peroxide or chlorhexidine gluconate mouthwashes, for reduction of SARS-CoV-2 viral load, and thus indicate a potential use for these products in the reduction of infectious risk associated with aerosol generating dental procedures and for SARS-CoV-2 infection control more generally. Our evidence supports inclusion of several of these mouthwashes into a randomised controlled trial to evaluate their efficacy and substantivity against SARS-CoV-2 *in-vivo*.

## Acknowledgements

The views expressed in this article are those of the authors and are not necessarily those of Public Health England, the National Health Service or the Department of Health and Social Care.

